# Characterization and transcriptome analysis reveal abnormal pollen germination in cytoplasmic male sterile tomato

**DOI:** 10.1101/2025.04.19.649671

**Authors:** Kosuke Kuwabara, Tohru Ariizumi

## Abstract

Cytoplasmic male sterility (CMS) is a phenomenon resulting from mitochondrial-nuclear incompatibility that is widely used in hybrid breeding. Although the mitochondrial gene *orf137* has been identified as the causal CMS gene in tomatoes, its function remains unclear. In this study, we characterized the male-sterile phenotypes and analyzed the CMS pollen transcriptome. Microscopic and calcium imaging analyses revealed that CMS pollen exhibited abnormal germination from multiple apertures, accompanied by elevated calcium concentrations and vesicle accumulation, which are typically observed in pollen tube tips. RNA-Seq analysis revealed 440 differentially expressed genes, including four *pectin methylesterase inhibitor (PMEI*) genes that were highly expressed in the pollen. PME activity was significantly reduced in CMS pollen, suggesting its association with abnormal pollen germination. ATP and reactive oxygen species (ROS) levels, which are key mediators of mitochondrial retrograde signaling (MRS), remained unchanged in CMS pollen, and the expression of the mitochondrial stress marker *AOX1*a was not elevated. These findings suggest that *orf137* triggers an alternative MRS pathway independent of ATP or ROS, potentially leading to *PMEI* upregulation and abnormal pollen germination. Our results reveal a previously unrecognized mechanism of CMS-induced male sterility in tomatoes involving nuclear gene regulation through unconventional mitochondrial signaling.

## 1. Introduction

Cytoplasmic male sterility (CMS) is caused by the incompatibility between the mitochondrial and nuclear genomes, leading to the abnormal development or function of male reproductive organs. CMS has been identified in more than 150 species of higher plants [1]. CMS plants cannot produce seeds through self-pollination; therefore, they are widely used in hybrid breeding systems because they eliminate manual emasculation and ensure high hybrid seed purity.

CMS is typically attributed to specific genes encoded by the mitochondrial genome, commonly known as CMS-causing genes. Different CMS genes have been identified across various plant species, and several models have been proposed to elucidate their mechanisms of action. Currently, four major models are widely discussed: (1) energy deficiency, (2) aberrant programmed cell death (PCD), (3) cytotoxicity, and (4) retrograde regulation models [2,3]. In the energy deficiency model, most CMS-causing gene products contain one or more transmembrane domains that can disrupt mitochondrial respiration by interfering with the components of the electron transport chain. This may reduce ATP synthesis and overproduction of reactive oxygen species (ROS). Since male reproductive development is highly dependent on ATP, a decline in mitochondrial ATP production is conjectured to directly disrupt the male reproductive organs [2,4]. In the aberrant PCD model, the tapetum of the anther undergoes precisely regulated PCD at specific developmental stages through a process mediated, in part, by ROS. The tapetum is crucial for supplying proteins, lipids, and other components essential for pollen development. However, excessive ROS production by CMS-causing proteins can trigger premature tapetum degeneration, leading to male sterility [5]. The cytotoxicity model has shown that a few CMS-inducing proteins exhibit cytotoxicity. For example, certain CMS proteins localize to the inner mitochondrial membrane, forming polymers or channels that cause electrolyte leakage and ultimately exert cytotoxic effects. These effects have been observed in *Escherichia coli* and yeast [4]. The retrograde regulation model hypothesizes that a few CMS-causing genes influence the expression of nuclear genes through mitochondrial retrograde signaling (MRS). Direct evidence of MRS involvement in CMS has been reported in Chinese wild (CW)-type rice. In this system, the *Rf17* gene, which encodes a protein with an acyl carrier protein synthase-like domain, restores fertility when a specific nucleotide substitution is present in the promoter region. In the cytoplasmic background of CMS, this substitution reduces *Rf17* expression, paradoxically leading to fertility restoration [6,7]. Recently, retrograde regulation has received increasing attention as a potential mechanism of CMS. Transcriptome analyses are commonly used to comprehensively identify nuclear genes whose expression is altered in the cytoplasmic background of CMS, with the goal of elucidating the downstream targets of mitochondrial signaling that contribute to male sterility.

Recently, our group identified *orf137* as a gene in tomato CMS lines but not in fertile cultivars [8]. This gene shares mild sequence similarity with *orf507*, a known CMS-causing gene in pepper [9]. Functional analysis using mitoTALEN-mediated knockout of *orf137* fully restored fertility in tomato CMS lines, providing strong evidence that *orf137* is the causal gene responsible for CMS in tomatoes [10]. However, the molecular mechanism of *orf137-*induced male sterility remains unclear. Therefore, we first investigated the phenotypic characteristics of male sterility in the tomato CMS line. We focused on detailed observations of vesicle and Ca^2+^ localization, both of which are critical for pollen germination, to better understand the abnormalities in this process. Additionally, we conducted transcriptome analysis of pollen to identify nuclear genes whose expression was altered in the CMS line and may be associated with male sterility. Based on these findings, we discuss the potential mechanistic relationship between *orf137*, changes in nuclear gene expression, and the manifestation of male sterility in tomatoes.

## 2. Results

### 2.1. Abnormal phenotypes during the pollen germination stage in the tomato CMS line

First, the development of anthers in tomato CMS lines “CMS[P]” was compared with the maintainer line “P.” No structural abnormalities were detected in the anthers of “CMS[P]” during their developmental stages. The tapetum layer, which plays an important role in pollen development, began to decompose from the tetrad stage (5 mm bud) and was completely degraded by the mature stage in the maintainer line. Similar degradation of the tapetum was observed in “CMS[P]” (Figure S1). Pollen staining experiments were performed. Before maturation, pollen accumulates starch, which is digested during anthesis [6]. Pollen stained with I_2_-KI solution was classified into three types: starch-digested, starch-engorged, and abnormal pollen. The proportion of each pollen type did not differ between the maintenance and CMS lines (Figure S2A, B). Staining with 4’,6-diamidino-2-phenylindole (DAPI) revealed similar results, with both the vegetative nucleus and sperm cells stained in both lines (Figure S2C). Transmission electron microscopy (TEM) observations of the internal structure of pollen showed organelles such as mitochondria, starch granules, and vacuoles, as well as the pollen wall, intine, and exine in both lines (Figure S3). These findings indicate that the anthers and pollen of “CMS[P]” develop normally, similar to those in the maintainer line.

We previously developed the tomato CMS line Dwarf “CMS[P]” by backcrossing the “CMS[P]” with the dwarf tomato cultivar “Micro-Tom.” This line retains the same mitochondrial genome as “CMS[P],” including the CMS-causing gene *orf137* [10]. The morphology of the plant was identical to that of “Micro-Tom” (Figure S4). The germination capability of pollen from “Micro-Tom” and Dwarf “CMS[P] was compared.” Pollen from “Micro-Tom” successfully germinated on both the artificial medium and the stigma. Conversely, pollen from Dwarf “CMS[P]” exhibited abnormal germination phenotypes, characterized by swelling or bursting at two or three apertures (pollen tube emergence sites). We quantified the proportion of pollen exhibiting these phenotypes at 2, 4, 6, and 24 h after incubation. In the “Micro-Tom”, approximately 90% of the pollen germinated within 2 h. In contrast, Dwarf “CMS[P]” pollen did not germinate (0% germination) within 2 to 24 h. The “Micro-Tom” displayed approximately 0% multiple swollen or burst apertures throughout the incubation period. In contrast, Dwarf “CMS[P]” exhibited 34% of the pollen with multiple swollen apertures at 2 h, which gradually decreased as incubation continued. The percentage of pollen grains with burst apertures increased from 9% at 2 h to 88% at 24 h (Figure 1). These findings indicate that pollen from tomato CMS lines undergoes abnormal morphological changes during germination induction, particularly swelling and bursting at multiple apertures.

**Figure 1.**
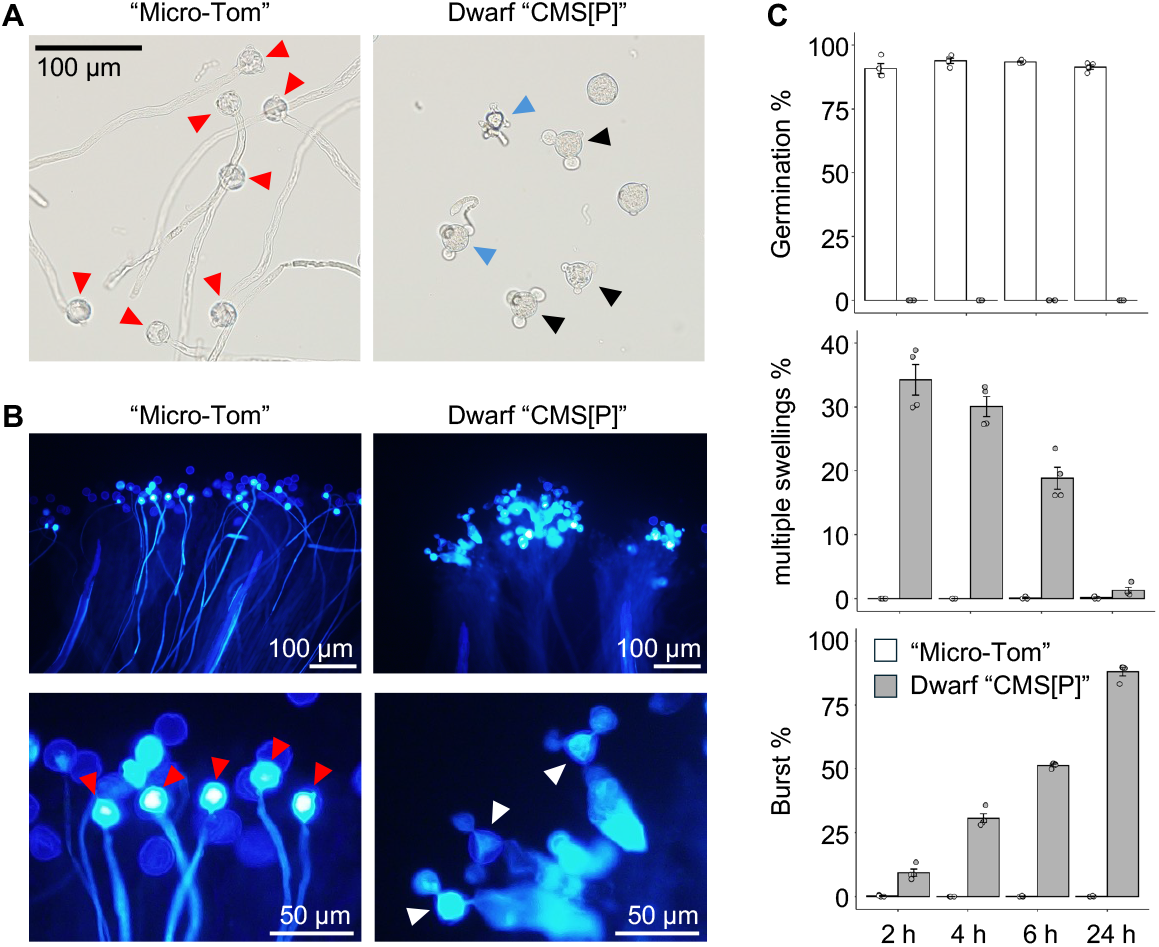
Abnormal pollen phenotypes in Dwarf “CMS[P].” (A) Pollen phenotypes after 4 h of incubation in liquid germination medium. Red arrowheads indicate germinated pollen in the “Micro-Tom”; black and blue arrowheads indicate pollen with multiple swollen apertures and burst apertures, respectively, in the Dwarf “CMS[P]”. (B) Morphology of pollen on the stigma 24 h after pollination. Red arrowheads indicate germinated pollen in the “Micro-Tom”; white arrowheads indicate pollen with multiple swollen or burst apertures. (C) Percentage of germinated pollen, pollen with multiple swollen apertures, and pollen with burst aperture. Data were obtained from four independent experiments (n = 4), and more than 300 pollen grains were observed in each experiment.

### 2.2. Pollen germination at multiple sites in tomato CMS line

The internal structure of pollen from Dwarf “CMS[P]” was investigated using TEM to explore its unusual pollen morphology following the initiation of pollen germination. In the pollen tube of “Micro-Tom,” vesicles were predominantly localized at the tip, whereas mitochondria and other organelles were absent. This observation was consistent across the six independent pollen samples (Figure 2A, B). These results are consistent with previously reported vesicle localization patterns in pollen tubes [11]. Similarly, numerous vesicles were observed in the swelling apertures of Dwarf “CMS[P],” with consistent results across all 11 independent pollens (Figure 2C, D).

**Figure 2.**
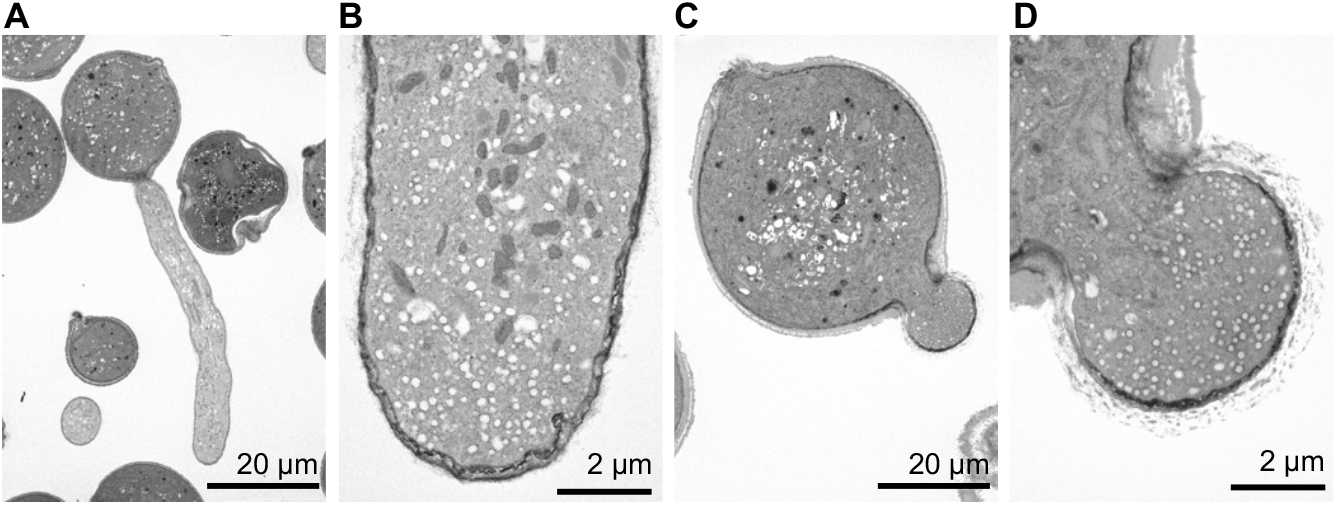
Transmission electron microscopy (TEM) of incubated pollen in the “Micro-Tom” and Dwarf “CMS[P]”. (A, B) TEM images of pollen structure in “Micro-Tom” and (C, D) Dwarf “CMS[P].” (B, D) Close-up images of the pollen tube tip and swollen aperture, respectively.

A cytosolic Ca^2+^ gradient is formed during pollen germination following hydration, and Ca^2+^ concentration ([Ca^2+^]_cyt_) increases at the site where the pollen tube emerges. Furthermore, a persistent Ca^2+^ gradient is maintained at the tip of the elongating pollen tube throughout its growth [12,13]. To visualize Ca^2+^ dynamics during pollen germination and pollen tube elongation, we used the Ca^2+^ indicator G-CaMP5 [13], which is specifically expressed in pollen under the control of the *Lat52* promoter. In “Micro-Tom” pollen, [Ca^2+^]_cyt_ initially increased at multiple germination apertures, but just before germination, a strong [Ca^2+^]_cyt_ signal was concentrated at a single aperture from which the pollen tube emerged. After germination, a strong [Ca^2+^]_cyt_ signal was consistently maintained at the tips of the elongated pollen tubes (Figure 3A, B). In Dwarf “CMS[P]” pollen, multiple apertures also showed strong [Ca^2+^]_cyt_ signals. Swelling of the germination apertures was observed, along with intense [Ca^2+^]_cyt_ accumulation, followed by similar swelling events in other apertures. Strong [Ca^2+^]_cyt_ signals persisted at the tips of the swollen apertures (Figure 3C). These results suggest that, rather than merely swelling after germination induction, Dwarf “CMS[P]” pollen attempts germination from multiple apertures, with [Ca^2+^]_cyt_ dynamics similar to those observed during normal pollen tube formation.

**Figure 3.**
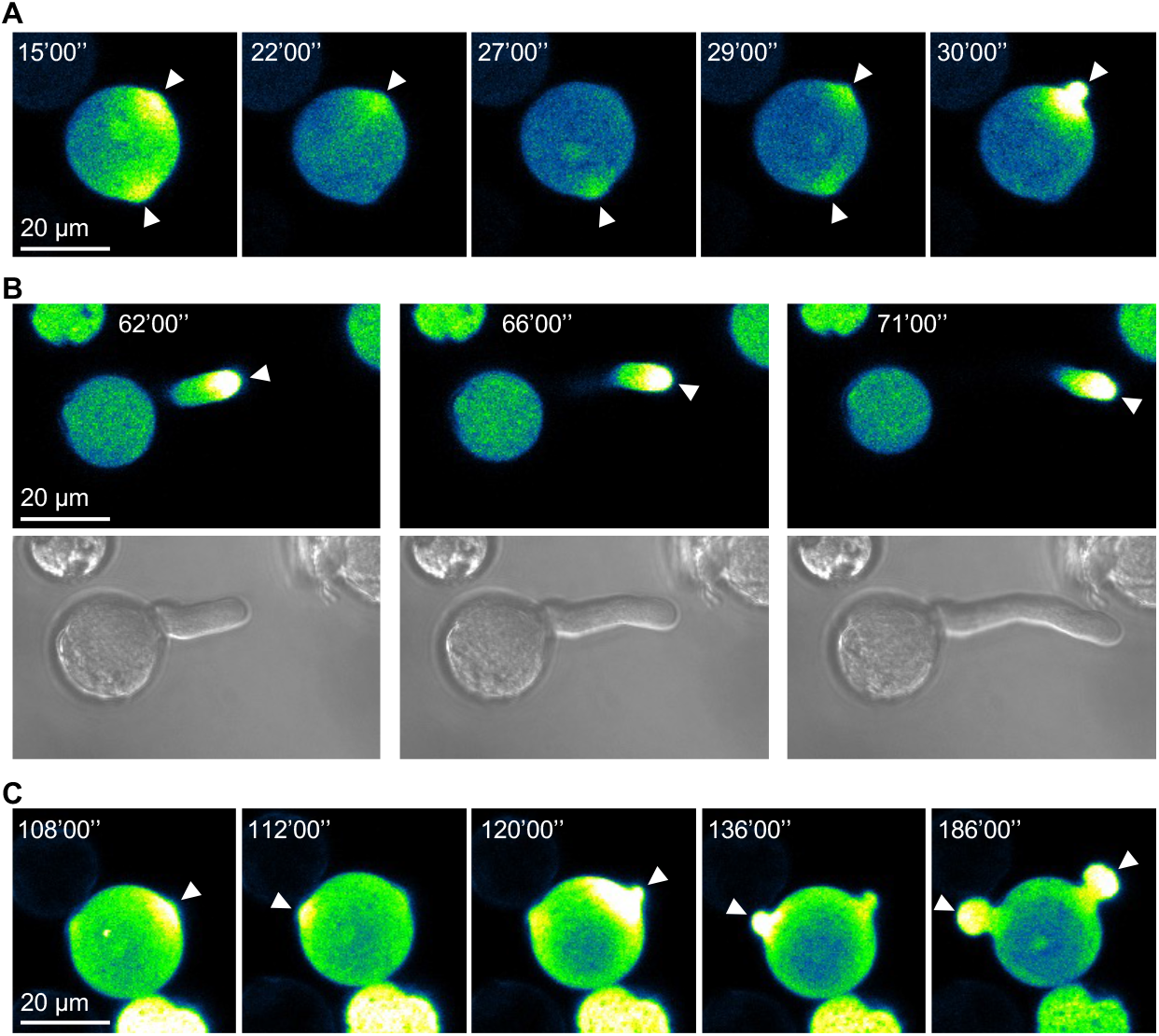
Time-course Ca^2+^ imaging of pollen germination and tube growth in “Micro-Tom” and aperture swelling in Dwarf “CMS[P].” (A) Time-lapse Ca^2+^ dynamics during pollen germination in “Micro-Tom.” (B) Time-lapse Ca^2+^ dynamics during pollen tube elongation in “Micro-Tom” (top: G-CaMP5 signal; bottom: bright-field image). (C) Time-lapse Ca^2+^ dynamics in Dwarf “CMS[P]” pollen showing aperture swelling. White arrowheads indicate regions with elevated Ca^2+^ signals.

### 2.3. Transcriptome analysis in pollen of tomato CMS line

“Micro-Tom” pollen begins to germinate after 15 min of incubation in a pollen germination medium (data not shown), and transcriptomic differences due to morphological changes may arise beyond this time point. To minimize the spatial and structural effects on gene expression, RNA-Seq was performed using pollen samples incubated for 10 min. As a result, a total of 440 differentially expressed genes (DEGs) were identified between “Micro-Tom” and Dwarf “CMS[P]” (|log_2_ fold change (FC|) > 1, false discovery rate (FDR) < 0.1), of which 177 were upregulated and 263 were downregulated in Dwarf “CMS[P]” compared to “Micro-Tom” (Figure 4).

**Figure 4.**
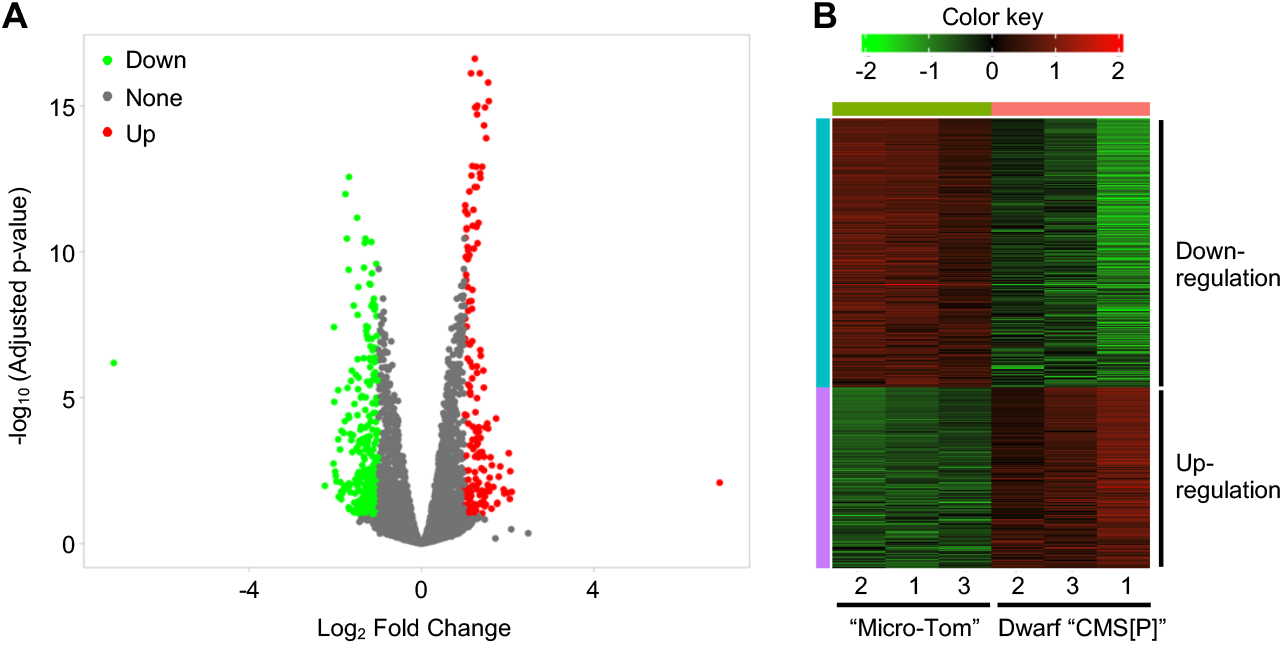
Transcriptome analysis of pollen from “Micro-Tom” and Dwarf “CMS[P]”. (A) Volcano plot of differentially expressed genes (DEGs) identified between “Micro-Tom” and Dwarf “CMS[P]” pollen. Genes with |log_2_ fold change| > 1 and false discovery rate (FDR) < 0.1 were considered DEGs. Red and green dots indicate significantly upregulated and downregulated genes in “CMS[P]” pollen, respectively, while gray dots represent non-significant genes. (B) Hierarchical heatmap clustering of DEGs between “Micro-Tom” and Dwarf “CMS[P].” TPM values were normalized using Z-score scaling. Each column represents a biological replicate (n = 3).

Among the DEGs, we focused on genes related to pectin modification because the pollen germination phenotype observed in Dwarf “CMS[P]” closely resembled that of the Arabidopsis *pme48* mutant, which exhibits pollen tube emergence or swelling from multiple apertures (or called “double-tipped”) [14]. Pectin methylesterase (PME) and PME inhibitor (PMEI) are key regulators of pectin methylation and have been reported to affect pollen tube formation [15]. To comprehensively identify *PME* and *PMEI* genes in tomatoes, we performed HMMER searches [16] using the Pfam domain profiles, Pfam01095 and Pfam04043, which correspond to *PME* and *PMEI*, respectively. Consequently, 77 *PME* and 48 *PMEI* genes were identified in the tomato nuclear genome. While none of the 77 *PME* genes showed significant expression changes, four of the 48 *PMEI* genes (*Solyc02g069300, Solyc05g005030, Solyc09g092350*, and *Solyc10g081670*) were significantly upregulated in Dwarf “CMS[P]” (Table S1). Subsequently, the tissue-specific expression patterns of all 48 *PMEI* genes were investigated using z-score normalization. Among them, five *PMEI* genes, including four identified as DEGs, were expressed in the male reproductive organs (Figure 5A). When examining the actual transcripts per million (TPM) values, the four *PMEI* DEGs were expressed at higher levels in incubated pollen than in the mature anthers. In contrast, *Solyc01g059940*, which was not identified as a DEG, exhibited similar expression levels between mature anthers and incubated pollen and the lowest expression among the five DEGs in the incubated pollen (Figure 5B). These findings suggest that the four *PMEI* DEGs are preferentially and highly expressed in pollen among the 48 *PMEI* genes and are likely responsible for regulating pectin methylesterification in pollen.

**Figure 5.**
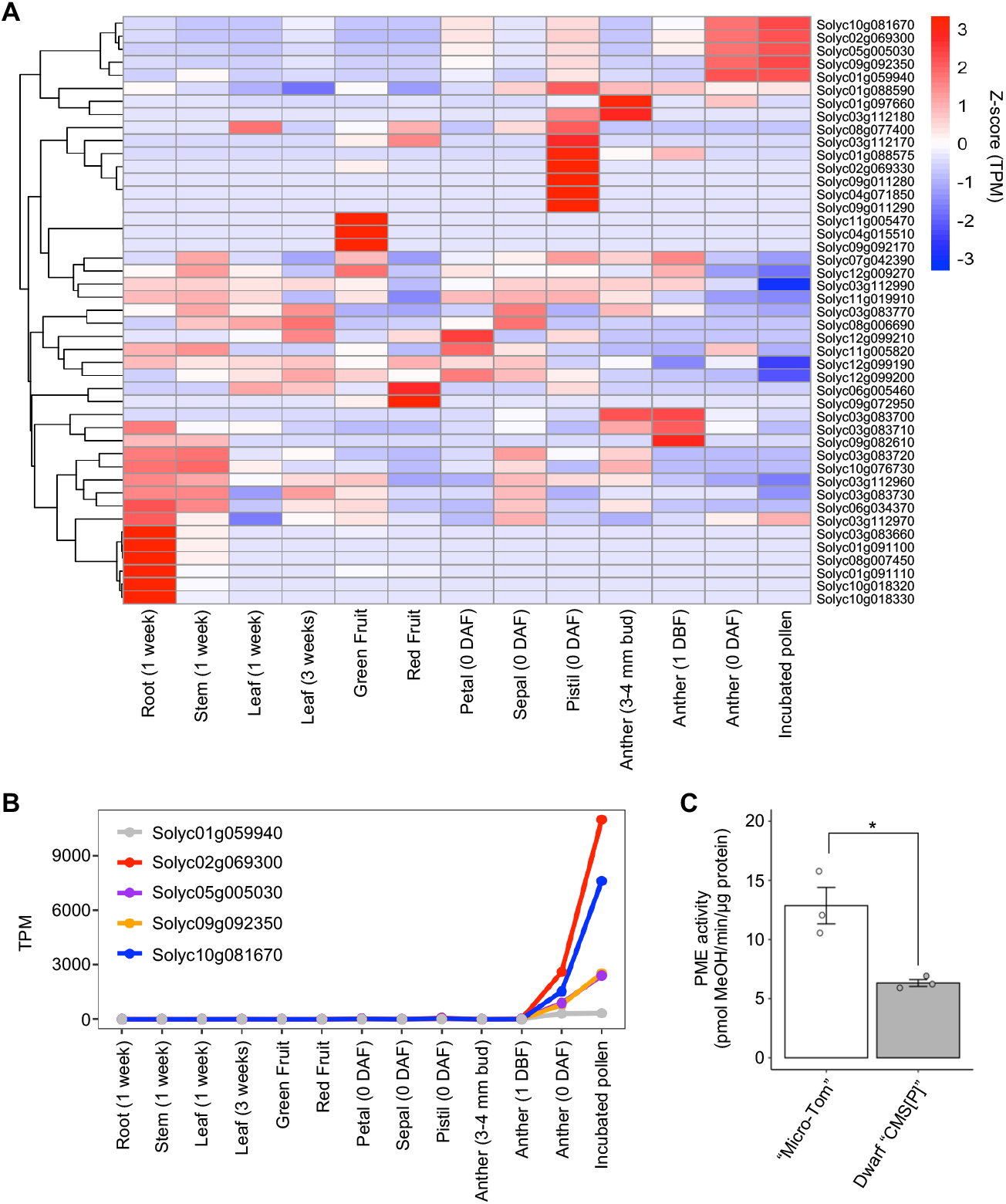
Expression profiles of *pectin methylesterase* (*PME*) and *PME inhibitor* (*PMEI*) genes and their activity in tomato male organs. (A) Heatmap showing tissue-specific expression patterns of 45 *PMEI* genes across various tomato tissues. Transcript per million (TPM) values were normalized using the Z-score. Three *PMEI* genes (Solyc08g016100, Solyc08g016110, and Solyc08g016120) were excluded from the analysis because their TPM values were zero in all tissues. (B) Line graph of TPM values for five highly expressed *PMEI* genes in male reproductive organs across different tissues. (C) PME enzymatic activity in the pollen of “Micro-Tom” and Dwarf “CMS[P].” Error bars indicate standard deviation (n = 3). Statistical significance was determined using the Student’s t-test. *P < 0.05.

To further assess the functional relevance of *PMEI* upregulation, PME enzyme activity was measured using total protein extracted from pollen incubated for 10 min. Consistent with the transcriptomic findings, PME activity in Dwarf “CMS[P]” pollen was significantly reduced to approximately half the level observed in “Micro-Tom” (Figure 5C). This reduction in PME activity indicates that the upregulated *PMEI* genes in Dwarf “CMS[P]” actively inhibit PME function. Overall, these results suggest that the elevated expression of specific *PMEI* genes and the consequent suppression of PME activity may contribute to the abnormal pollen germination phenotype characteristic of Dwarf “CMS[P],” particularly the emergence from multiple apertures.

### 2.4. Assessment of ATP and ROS production in tomato CMS strain pollen

All four *PMEI* genes that were strongly expressed in pollen were upregulated in Dwarf “CMS[P],” suggesting the existence of a regulatory mechanism in which the mitochondrial gene *orf137* influences the expression of these nuclear genes via mitochondrial retrograde signaling (MRS). ATP and ROS are well-known inducers of MRS [17]. In other plant species, CMS-causing genes induce a loss of function in the respiratory chain complex, resulting in decreased ATP and increased ROS levels in male organs [2,4]. Therefore, we investigated whether ATP and ROS production fluctuated in the pollen of tomato CMS lines. Fluorescent labeling of ROS within the pollen using H_2_DCFDA revealed comparable fluorescence and intensities in the pollen of “Micro-Tom” and Dwarf “CMS [P]” (Figure 6A, B). In addition, ATP content was similar in the two lines (Figure 6C). These results suggest that pollen from tomato CMS lines maintains normal ATP and ROS levels. *AOX1a* is commonly used as a marker gene for mitochondrial stress responsiveness because its expression increases during stress [17,18]. RNA-Seq data showed no variation in *AOX1a* expression levels between “Micro-Tom” and Dwarf “CMS[P]” (Figure 6D). These observations imply that ATP production and ROS development are normal in Dwarf “CMS [P]” pollen and that there are distinct signaling pathways from conventional mitochondrial stress that regulate the four nuclear-encoded *PMEI* genes.

**Figure 6.**
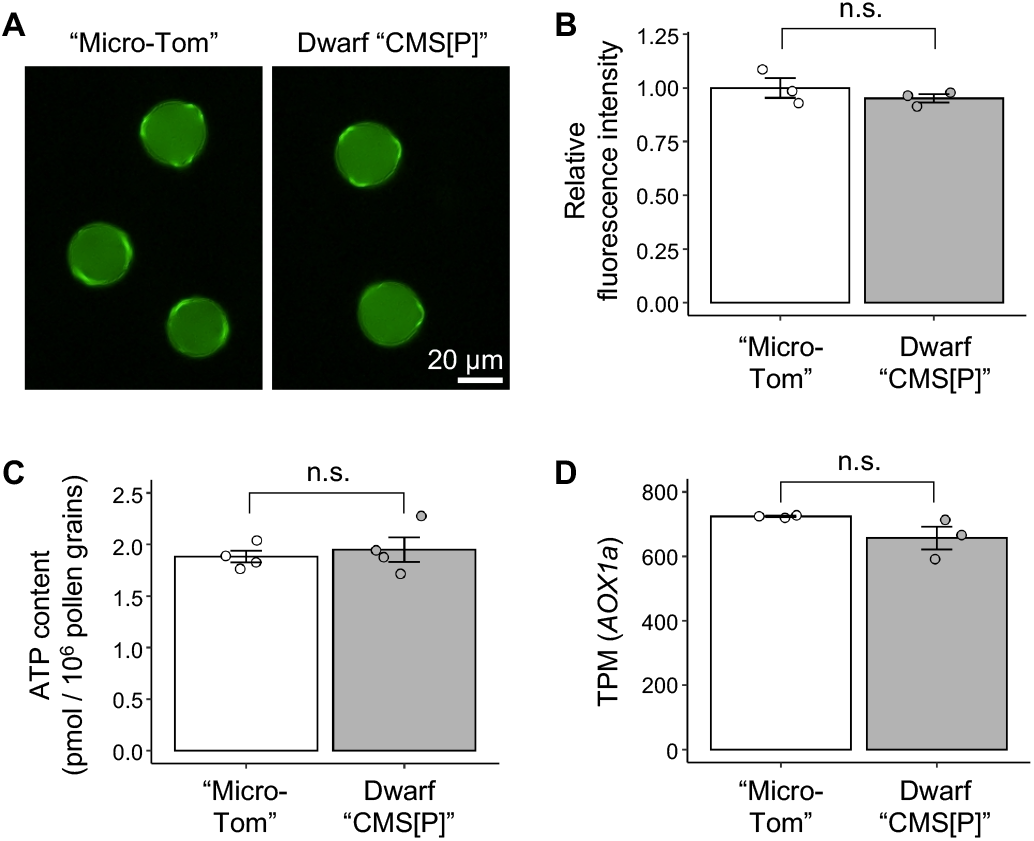
ROS levels, ATP content, and *AOX1a* expression in pollen. (A) Representative H_2_DCFDA-stained fluorescent images of pollen showing reactive oxygen species (ROS) in “Micro-Tom” and Dwarf “CMS[P].” (B) Quantification of fluorescence intensity in (A). Data are shown as mean ± SE from three independent biological replicates (n = 3). More than 100 pollen grains were measured for each replicate. (C) ATP content in pollen, measured as pmol per 10^6^ pollen grains. n =3. (D) Expression levels of the *AOX1a* gene in pollen as transcripts per million (TPM) values. No statistically significant difference (n.s.) was observed between “Micro-Tom” and Dwarf “CMS[P].”

## 3. Discussion

### 3.1. Tomato CMS lines show exhibit an unusual male sterility phenotype

In a previous study, we identified *orf137* as the gene responsible for CMS in tomatoes [10]. In this study, we investigated the mechanism by which *orf137* induces male sterility in tomatoes. The plant structure and floral organs showed no morphological abnormalities, and the anthers appeared normal. The premature degradation of the anther’s tapetal layer has been reported in many CMS lines [2], which is believed to result from the excessive production of ROS by the protein encoded by the CMS-causing gene. However, in the tomato CMS line, tapetum degradation commenced at a time comparable to that in the maintainer line and progressed normally. These findings indicate that the anthers in the tomato CMS line are functionally normal and that *orf137* does not induce defects in sporophytic male organs. Additionally, pollen staining showed no discernible differences between the CMS and maintainer lines, suggesting that pollen development proceeded normally in the CMS lines. The only phenotypic difference observed was in the pollen germination. The maintainer line’s pollen germinated normally on both the stigma and germination medium, whereas the CMS line’s pollen exhibited an abnormal phenotype characterized by swelling or bursting at multiple apertures.

TEM revealed concentrated vesicles at the tip of the pollen tubes in “Micro Tom.” In contrast, mitochondria and other organelles were not observed at the tips of the tubes. This phenomenon is well documented in Arabidopsis pollen, where vesicles are actively transported to the pollen tube tip along actin filaments by myosin motors. In contrast, larger organelles, such as Golgi bodies and mitochondria, move along actin filaments in the shank region of the pollen tube rather than accumulating at the tip [11,19]. A similar phenomenon was observed in the swollen apertures of CMS pollen grains. In addition, it is well known that [Ca^2+^]_cyt_ accumulates at the site where the pollen tube emerges during germination, and a consistent Ca^2+^ gradient is maintained at the tip of the pollen tube [12,13]. This tip-focused Ca^2+^ gradient is important for controlling cell growth, vesicular transport, and intracellular signaling [20]. These phenomena were observed in the pollen tubes of “Micro-Tom” and the swollen apertures of Dwarf “CMS[P]” pollen. Overall, TEM and Ca ^2+^ imaging results indicated that phenomena consistent with pollen germination occurred at multiple apertures in tomato CMS pollen grains. These findings suggest that tomato CMS pollen exhibits a unique phenotype characterized by germination from multiple apertures.

### 3.2. Upregulated PMEI gene expression is associated with male sterility in the tomato CMS lines

PME catalyzes the demethylesterification of homogalacturonan, which is a major pectin component. PME activity is precisely regulated by PMEI through direct protein–protein interactions. The balance between PME and PMEI determines the degree of pectin methylesterification, which affects the mechanical properties of the cell walls. Mutations in *PME* or *PMEI* often result in various phenotypic changes, including altered plant growth, responses to abiotic stress, and defects in pollen development. In Arabidopsis, the *pme48* mutant shows reduced PME activity in pollen and a higher frequency of pollen grains exhibiting swelling or emergence of pollen tube-like structures from two of the three apertures, a phenotype not typically observed in wild-type plants [14]. In the tomato CMS lines examined in this study, no significant changes in *PME* gene expression were observed. However, four *PMEI* genes that were highly expressed in mature pollen were upregulated. A reduction in PME enzymatic activity was consistently observed in the pollen of the CMS line, likely due to the increased expression of *PMEI* genes. Considering the relationship between reduced PME activity and abnormal pollen germination observed in the Arabidopsis *pme48* mutant [14], the decreased PME activity detected in tomato CMS pollen was conjectured to be associated with the emergence of pollen tubes from multiple apertures. However, the mechanism by which reduced PME activity results in pollen germination from multiple apertures remains unclear and requires further investigation.

### 3.3. Normal ATP and ROS production in tomato CMS line

Most CMS-causing genes encode proteins with one or more transmembrane domains and interfere with mitochondrial respiration by disrupting components of the electron transport chain. This disruption can result in decreased ATP production and elevated ROS levels [2–4]. The tomato CMS-causing gene *orf137* encodes a protein predicted to have a single transmembrane domain [8] and was initially hypothesized to reduce ATP synthesis and elevate ROS levels in the pollen. However, contrary to this hypothesis, ATP and ROS levels were comparable between the CMS and maintainer lines. Alterations in mitochondrial ATP and ROS levels are recognized as triggers for MRS, a process in which the functional state of mitochondria regulates the expression of nuclear genes [17]. CMS-causing genes have been proposed to activate MRS to mediate changes in nuclear gene expression [2]. For example, in maize, the CMS-causing gene *orf355* regulates the expression of the nuclear transcription factor *ZmDREB1.7* via an MRS-dependent pathway [18]. Direct evidence of MRS in rice CW-CMS was provided by the *Rf17* gene, which encodes a protein with an acyl carrier protein synthase-like domain. A single nucleotide substitution in the promoter region of *Rf17* led to reduced expression in the CMS background, paradoxically restoring fertility [6]. In many CMS systems, increased expression of mitochondrial stress marker genes, such as *AOX1a*, has been used as indirect evidence of MRS activation. *AOX1a* upregulation has been reported in maize CMS-S and rice CW-CMS lines [6,18]. In contrast, our RNA-Seq data revealed no change in *AOX1a* expression between “Micro-Tom” and Dwarf “CMS[P]” pollen, which is consistent with the unaltered ATP and ROS levels in the CMS line. Despite the lack of ATP or ROS fluctuations, we observed upregulation of four nuclear-encoded *PMEI* genes in CMS pollen. This suggests that *orf137* may induce an alternative MRS pathway that is independent of ATP or ROS. Other known inducers of MRS include the NADH/NAD ^+^ ratio, intracellular pH, free Ca^2+^ concentration, and changes in the mitochondrial membrane potential [17].

In wheat CMS, which carries the cytoplasm of *Aegilops crassa*, the CMS-causing gene *orf260* has been suggested to cause the homeotic transformation of stamens into pistil-like organs via MRS-mediated alteration of *B-class MADS-box* genes. In this system, *Wheat Calmodulin-Binding Protein 1* (*WCBP1*) has been proposed as a component of a Ca^2+^-dependent signaling pathway, suggesting that *orf260* may initiate MRS through Ca^2+^ signaling [21,22]. These findings suggest that *orf137* in tomato may similarly activate an unconventional MRS pathway, potentially involving Ca^2+^ or other signaling molecules, leading to upregulation of *PMEI* genes. Future studies focusing on the dynamics of signaling pathways beyond ATP and ROS in CMS pollen are essential to elucidate the molecular connections between *orf137* and nuclear gene regulation.

## 4. Materials and Methods

### 4.1. Plant materials

Tomato CMS line “CMS[P]” was previously developed through an asymmetric cell fusion, where the tomato cultivar “P” served as the nuclear donor and the wild potato relative *Solanum acaule* as the cytoplasmic donor, followed by repeated backcrossing using “P” [23]. The plants were cultivated in greenhouses at the University of Tsukuba. The dwarf “CMS[P]” variant was developed from “CMS[P]” by backcrossing with the tomato dwarf cultivar “Micro-Tom” (TOMJPF0001) [24] in our previous study [8]. They were cultivated at 23 °C under a 16/8 h light/dark cycle.

### 4.2. Phenotypic analysis of anthers and pollen

Histological examination of the anthers was performed using a previously described method [25]. Pollen germination was assessed in a liquid germination medium and on the stigma, according to previously described methods [10]. For DAPI staining, pollen from the anthesis stage was fixed in ethanol: acetic acid 3:1. The pollen was then stained using DAPI solution, which contained 0.1 M phosphate buffer (pH 7.0), 0.5 M EDTA, Triton X-100, and 3.0 µg/mL DAPI, for 5 min and observed under ultraviolet light using an OLYMPUS BX53 microscope. For I_2_-KI staining, pollen was collected at the anthesis stage, it was stained with a 2 w/v% I_2_-KI solution. Pollen grains were classified as starch-digested, starch-engorged, or abnormal and counted under a light microscope. ROS levels in the pollen were measured as previously described [26]. The fluorescence intensity of individual pollen grains was quantified using the ImageJ software [27].

The pollen structure was examined using TEM. Pollen grains were incubated in a liquid germination medium for 1 h and then fixed using a two-step fixation protocol. First, samples were treated with 4% paraformaldehyde containing 10% sucrose and 15.1% polyethylene glycol 6000 in 0.05 M cacodylate buffer (pH 7.4) at 4°C for 1 h. This was followed by a second fixation using 2% paraformaldehyde and 2% glutaraldehyde in the same buffer at 4°C for 16 h. After three rinses with cacodylate buffer, the pollen was post-fixed with 2% osmium tetroxide in cacodylate buffer overnight at 4°C. Dehydration was achieved using a graded ethanol series (50%, 70%, 90%, and 100% ethanol) at 25°C. To ensure complete dehydration, the samples were kept in 100% ethanol for two days. Subsequently, the samples were infiltrated with propylene oxide and a 1:1 mixture of propylene oxide and epoxy resin (Quetol-651, Nisshin) for 6 h and finally embedded in pure resin overnight. Polymerization was performed at 60°C for 48 h. Ultrathin sections (80 nm) were prepared using an ultramicrotome (Leica Ultracut UCT, Wetzlar, Germany) and stained with 2% uranyl acetate and lead citrate. The sections were examined under a TEM (JEM-1400 Plus, JEOL) at 100 kV, and images were captured using a CCD camera (EM-14830RUBY2, JEOL).

### 4.3. Ca^2+^ imaging in pollen

To create a plasmid that expressed the Ca^2+^ probe in pollen, the G-CaMP5 sequence was amplified from pCMV-GCaMP5G (Addgene #31788) using the following primers (5’-GGTTTAGTGAACCGTCAGA-3’ and 5’-AGGAGAGTTGTTGATTCACTTCGCTGTCATCATTTG-3’). The *Lat52* promoter fragment was amplified from the genomic DNA of “Micro-Tom” using primers (5’-AGACCAAAGGGCAATATACTCGACTCAGAAGGTATTG-3’ and 5’-ACGGTTCACTAAACCTTTAAATTGGAATTTTTTTTTTTGGTGT-3’). The vector backbone was amplified from the pBI121 plasmid (Clontech) using primers (5’-ATCAACAACTCTCCTGGC-3’ and 5’-ATTGCCCTTTGGTCTTCT-3’). These fragments were ligated using the In-Fusion HD Cloning Kit (Takara Bio) to construct the *Lat52::G-CaMP5* vector. The resulting vector was introduced into *Agrobacterium tumefaciens* strain GV3101. Transformation of the tomato cultivar “Micro-Tom” was performed according to the previously described method [28], and it was cross-pollinated with Dwarf “CMS[P].”

Pollen was collected from plants expressing *Lat52::G-CaMP5* and placed on a solid germination medium (12% w/v sucrose, 1.2 mM H_3_BO_3_, 1.6 mM Ca(NO_3_)_2_, 1 mM MgSO_4_, 0.1 mM K_2_HPO_4_, 0.5 w/v% agarose). The solid medium was inverted in a glass-bottom dish for Ca^2+^ imaging, which was performed using a confocal laser scanning microscope (LSM700; Zeiss). The G-CaMP5 signal was detected by excitation at 488 nm, and detection was performed between 490 and 635 nm.

### 4.4. RNA expression analyses

Total RNA was extracted from pollen that had been incubated for 10 min in a liquid germination medium using the RNeasy Plant Mini Kit (Qiagen). The extracted RNA was treated with RNase-free DNase (Qiagen) and used for the preparation of a sequence library using the NEBNext® Ultra™ Directional RNA Library Prep Kit for Illumina, with poly(A) selection and strand-specific protocol. Sequencing was performed on a NovaSeq 6000 platform (Illumina) to produce 150 bp paired-end reads, generating approximately 2 Gb and 13.3 million reads per sample. In addition, RNA-seq data from various tomato tissues [29], as detailed in Supplementary Table 2, were retrieved using Fasterq-dump (https://github.com/ncbi/sra-tools/wiki/01.-Downloading-SRA-Toolkit).

Trim_galore (https://github.com/FelixKrueger/TrimGalore) was used to trim adaptors and low-quality reads, with option q 30. Read mapping was performed using HISAT2 [30] with the nuclear genome of the cultivated tomato cv. Heinz 1706 (SL4.0) [31] as the reference genome. Transcript quantification was performed using TPMCalculator [32]. The resulting count data were uploaded to the integrated Differential Expression and Pathway analysis) (iDEP2.0) [33] for normalization and identification of differentially expressed genes (DEGs) using DESeq2 [34]. Genes with |log_2_ Fold Change| > 1 and FDR < 0.1 were considered significantly differentially expressed. To comprehensively identify *PME* and *PMEI* genes in tomatoes, HMMER searches [16] were performed using the Pfam domain profiles, Pfam01095 and Pfam04043, which correspond to *PME* and *PMEI*, respectively.

### 4.5. PME activity

PME activity was measured as previously described [35]. Fifty anthers at anthesis were collected and vortexed in 5 mL of liquid pollen germination medium to release pollen. The suspension was filtered through a 30 µm filter (CellTrics; Sysmex) and shaken at room temperature for 10 min. Pollen was pelleted by centrifugation at 1,000 *×g* for 1 min at 4°C. The resulting pellet was resuspended in 100 µL of protein extraction buffer (1 M NaCl, 200 mM Na_2_HPO_4_, pH 6.2) supplemented with 1 µL of Protease Inhibitor Cocktail (Sigma-Aldrich), and the pollen was disrupted using a plastic pestle. Following centrifugation at 15,000 *×g* for 15 min at 4°C, the supernatant was collected. Protein concentration was measured using the Bradford assay [36], and the concentration was adjusted to 2 µg/µL using the protein extraction buffer. Subsequently, 8 µL of the protein solution was mixed with 72 µL of 20 mM Na_2_HPO_4_ (pH 6.2), and 200 µL of 0.5% pectin (Sigma-Aldrich) was added. The mixture was incubated at 30°C for 3 h and then heat-inactivated at 100°C for 10 min. After cooling, 70 µL of the reaction solution was transferred to a 96-well plate, and 30 µL of 0.001 U/µL alcohol oxidase (Sigma-Aldrich) was added to each well. The plate was shaken for 15 min, and then 100 µL of reaction buffer (0.02 M acetylacetone, 2 M ammonium acetate, 0.05 M acetic acid) was added and shaken for 30 s. The plates were incubated at 68°C for 10 min and then cooled to 4°C for 10 min. The absorbance at 412 nm was measured using a microplate reader (SpectraMax M2; Molecular Devices). A standard curve was prepared using a methanol dilution series (3.5–70 nmol) treated in the same manner.

### 4.6. ATP quantification assay in pollen

ATP quantification was performed as previously described [37]. Anthers from ten flowers at the anthesis stage were collected in a pollen germination medium and vortexed to release the pollen. The suspension was passed through a 30 µm filter (CellTrics; Sysmex). A 100 µL aliquot of the pollen suspension was added to a 1.5 mL tube containing 400 µL of 7% perchloric acid and 10 mM EDTA and placed on ice for 15 min. Then, 100 µL of 5 N KOH and 1 M triethanolamine were added, and the mixture was cooled on ice for 10 min. Following centrifugation at 10,000 *×g* for 5 min at 4°C to precipitate the neutralized salts, 10 µL of the supernatant was mixed with 990 µL of 50 mM Tris-HCl buffer (pH 8.0). Then, 100 µL of the diluted supernatant was transferred to a white 96-well plate, and 100 µL of the ENLITEN rLuciferase/Luciferin Reagent (Promega) was added. The solution was gently mixed by pipetting, and luminescence was measured using a Fluoroskan Ascent FL luminometer (Thermo Fisher Scientific) with an integration time of 10 s. A standard curve was prepared using a 10-fold serial dilution of 10^−7^ M ATP standard solution (Promega) in 50 mM Tris-HCl buffer (pH 8), ranging from 10^−7^ to 10^−11^ M. As a blank control, the pollen germination medium without pollen was subjected to the same ATP extraction procedure described above. The pollen concentration in the suspension was determined using a cell counter (OneCell).

## Short legends for Supplementary Materials

**Table S1**. DEG analysis for *PME* and *PMEI* genes.

**Table S2**. RNA-Seq data used in this study.

**Figure S1**. Developmental progression of anther tissues in “P” and “CMS[P]”.

**Figure S2**. Pollen staining experiments in “P” and “CMS[P]”.

**Figure S3**. Transmission electron microscopy (TEM) analysis for pollen structure in “P” and “CMS[P]”.

**Figure S4**. Plant morphology of “Micro-Tom” and Dwarf “CMS[P]”.

## Author Contributions

K.K. and T.A. conceived and coordinated the study. K.K. performed the plant experiments. K.K. and T.A. performed sequencing analysis. K.K. wrote the manuscript.

## Funding

This research was funded by the Project of the NARO Bio-Oriented Technology Research Advancement Institution (Research Program on Development of Innovative Technology, Grant Number: JPJ007097) to T.A., the Japan Society for the Promotion of Science (JSPS) KAKENHI (Grant Number: 21H02181) to T.A., and the JSPS Research Fellowships for Young Researchers (Grant Number: 21J20479) to K.K.

## Institutional Review Board Statement

Not applicable.

## Informed Consent Statement

Not applicable.

## Data Availability Statement

The raw RNA-Seq sequences are available at the Sequence Read Archive (SRA) under the BioPro-ject ID PRJNA1250823.

## Acknowledgments

We thank all the technical and administrative members of the T-PIRC Center at the University of Tsukuba. Computations were partially performed on the NIG supercomputer at the ROIS National Institute of Genetics. Micro-Tom (TOMJPF0001) was provided by National BioResource Project Tomato (NBRP tomato).

## Conflicts of Interest

The authors declare no conflicts of interest.

## Abbreviations

The following abbreviations are used in this manuscript:

CMS: Cytoplasmic male sterility
ROS: Reactive oxygen species
MRS: Mitochondrial retrograde signaling
PCD: Programmed cell death
TEM: Transmission electron microscope
PME: Pectin methylesterase
PMEI: Pectin methylesterase inhibitor

## Notes

### Competing Interest Statement

The authors have declared no competing interest.

## References

1. Bohra, A.; Jha, U.C.; Adhimoolam, P.; Bisht, D.; Singh, N.P. Cytoplasmic Male Sterility (CMS) in Hybrid Breeding in Field Crops. Plant Cell Rep 2016, 35, 967–993, doi:10.1007/s00299-016-1949-3.

2. Chen, L.; Liu, Y.-G. Male Sterility and Fertility Restoration in Crops. Annu. Rev. Plant Biol. 2014, 65, 579– 606, doi:10.1146/annurev-arplant-050213-040119.

3. Bhattacharya, J.; Nitnavare, R.; Bhatnagar-Mathur, P.; Palakolanu, S.R. Cytoplasmic Male Sterility-Based Hybrids: Mechanistic Insights. Planta 2024, 260, doi:10.1007/s00425-024-04532-w.

4. Hu, J.; Huang, W.; Huang, Q.; Qin, X.; Yu, C.; Wang, L.; Li, S.; Zhu, R.; Zhu, Y. Mitochondria and Cytoplasmic Male Sterility in Plants. Mitochondrion 2014, 19, 282–288, doi:10.1016/j.mito.2014.02.008.

5. Biswas, R.; Chaudhuri, S. The Tale of Tapetum: From Anther Walls to Pollen Wall. Nucleus 2024, 67, 611– 630, doi:10.1007/s13237-024-00510-5.

6. Fujii, S.; Toriyama, K. Suppressed Expression of RETROGRADE-REGULATED MALE STERILITY Restores Pollen Fertility in Cytoplasmic Male Sterile Rice Plants. Proceedings of the National Academy of Sciences 2009, 106, 9513–9518, doi:10.1073/pnas.0901860106.

7. Suketomo, C.; Kazama, T.; Toriyama, K. Fertility Restoration of Chinese Wild Rice-Type Cytoplasmic Male Sterility by CRISPR/Cas9-Mediated Genome Editing of Nuclear-Encoded RETROGRADE-REGULATED MALE STERILITY. Plant Biotechnol (Tokyo) 2020, 37, 285–292, doi:10.5511/plantbiotechnology.20.0326b.

8. Kuwabara, K.; Harada, I.; Matsuzawa, Y.; Ariizumi, T.; Shirasawa, K. Organelle Genome Assembly Uncovers the Dynamic Genome Reorganization and Cytoplasmic Male Sterility Associated Genes in Tomato. Hortic Res 2021, 8, 250, doi:10.1038/s41438-021-00676-y.

9. Gulyas, G.; Shin, Y.; Kim, H.; Lee, J.-S.; Hirata, Y. Altered Transcript Reveals an Orf507 Sterility-Related Gene in Chili Pepper (Capsicum Annuum L.). Plant Mol Biol Rep 2010, 28, 605–612, doi:10.1007/s11105-010-0182-4.

10. Kuwabara, K.; Arimura, S.-I.; Shirasawa, K.; Ariizumi, T. Orf137 Triggers Cytoplasmic Male Sterility in Tomato. Plant Physiol 2022, 189, 465–468, doi:10.1093/plphys/kiac082.

11. Ruan, H.; Li, J.; Wang, T.; Ren, H. Secretory Vesicles Targeted to Plasma Membrane During Pollen Germination and Tube Growth. Front. Cell Dev. Biol. 2021, 8, doi:10.3389/fcell.2020.615447.

12. Iwano, M.; Shiba, H.; Miwa, T.; Che, F.-S.; Takayama, S.; Nagai, T.; Miyawaki, A.; Isogai, A. Ca2+ Dynamics in a Pollen Grain and Papilla Cell during Pollination of Arabidopsis. Plant Physiology 2004, 136, 3562–3571, doi:10.1104/pp.104.046961.

13. Diao, M.; Qu, X.; Huang, S. Calcium Imaging in Arabidopsis Pollen Cells Using G-CaMP5. Journal of Integrative Plant Biology 2018, 60, 897–906, doi:10.1111/jipb.12642.

14. Leroux, C.; Bouton, S.; Kiefer-Meyer, M.-C.; Fabrice, T.N.; Mareck, A.; Guénin, S.; Fournet, F.; Ringli, C.; Pelloux, J.; Driouich, A.; et al. PECTIN METHYLESTERASE48 Is Involved in Arabidopsis Pollen Grain Germination. Plant Physiology 2015, 167, 367–380, doi:10.1104/pp.114.250928.

15. Bosch, M.; Hepler, P.K. Pectin Methylesterases and Pectin Dynamics in Pollen Tubes. Plant Cell 2005, 17, 3219–3226, doi:10.1105/tpc.105.037473.

16. Finn, R.D.; Clements, J.; Eddy, S.R. HMMER Web Server: Interactive Sequence Similarity Searching. Nucleic Acids Res 2011, 39, W29–W37, doi:10.1093/nar/gkr367.

17. Khan, K.; Tran, H.C.; Mansuroglu, B.; Önsell, P.; Buratti, S.; Schwarzländer, M.; Costa, A.; Rasmusson, A.G.; Van Aken, O. Mitochondria-Derived Reactive Oxygen Species Are the Likely Primary Trigger of Mitochondrial Retrograde Signaling in Arabidopsis. Current Biology 2024, 34, 327-342.e4, doi:10.1016/j.cub.2023.12.005.

18. Xiao, S.; Zang, J.; Pei, Y.; Liu, J.; Liu, J.; Song, W.; Shi, Z.; Su, A.; Zhao, J.; Chen, H. Activation of Mitochondrial Orf355 Gene Expression by a Nuclear-Encoded DREB Transcription Factor Causes Cytoplasmic Male Sterility in Maize. Molecular Plant 2020, 13, 1270–1283, doi:10.1016/j.molp.2020.07.002.

19. Zhao, L.; Rehmani, M.S.; Wang, H. Exocytosis and Endocytosis: Yin-Yang Crosstalk for Sculpting a Dynamic Growing Pollen Tube Tip. Front. Plant Sci. 2020, 11, doi:10.3389/fpls.2020.572848.

20. Steinhorst, L.; Kudla, J. Calcium - a Central Regulator of Pollen Germination and Tube Growth. Biochimica et Biophysica Acta (BBA) - Molecular Cell Research 2013, 1833, 1573–1581, doi:10.1016/j.bbamcr.2012.10.009.

21. Yamamoto, M.; Shitsukawa, N.; Yamada, M.; Kato, K.; Takumi, S.; Kawaura, K.; Ogihara, Y.; Murai, K. Identification of a Novel Homolog for a Calmodulin-Binding Protein That Is Upregulated in Alloplasmic Wheat Showing Pistillody. Planta 2013, 237, 1001–1013, doi:10.1007/s00425-012-1812-x.

22. Hama, E.; Takumi, S.; Ogihara, Y.; Murai, K. Pistillody Is Caused by Alterations to the Class-B MADS-Box Gene Expression Pattern in Alloplasmic Wheats. Planta 2004, 218, 712–720, doi:10.1007/s00425-003-1157-6.

23. Melchers, G.; Mohri, Y.; Watanabe, K.; Wakabayashi, S.; Harada, K. One-Step Generation of Cytoplasmic Male Sterility by Fusion of Mitochondrial-Inactivated Tomato Protoplasts with Nuclear-Inactivated Solanum Protoplasts. Proceedings of the National Academy of Sciences 1992, 89, 6832–6836, doi:10.1073/pnas.89.15.6832.

24. Scott, J.W.; Station, U. of F.A.E.; Harbaugh, B.K. Micro-Tom: A Miniature Dwarf Tomato; Agricultural Experiment Station, Institute of Food and Agricultural Sciences, University of Florida, 1989;

25. Hao, S.; Ariizumi, T.; Ezura, H. SEXUAL STERILITY Is Essential for Both Male and Female Gametogenesis in Tomato. Plant and Cell Physiology 2017, 58, 22–34, doi:10.1093/pcp/pcw214.

26. Gao, X.-Q.; Liu, C.Z.; Li, D.D.; Zhao, T.T.; Li, F.; Jia, X.N.; Zhao, X.-Y.; Zhang, X.S. The Arabidopsis KINβγ Subunit of the SnRK1 Complex Regulates Pollen Hydration on the Stigma by Mediating the Level of Reactive Oxygen Species in Pollen. PLOS Genetics 2016, 12, e1006228, doi:10.1371/journal.pgen.1006228.

27. Abràmoff, M.D. Image Processing with ImageJ.

28. Sun, H.-J.; Uchii, S.; Watanabe, S.; Ezura, H. A Highly Efficient Transformation Protocol for Micro-Tom, a Model Cultivar for Tomato Functional Genomics. Plant and Cell Physiology 2006, 47, 426–431, doi:10.1093/pcp/pci251.

29. Ezura, K.; Ji-Seong, K.; Mori, K.; Suzuki, Y.; Kuhara, S.; Ariizumi, T.; Ezura, H. Genome-Wide Identification of Pistil-Specific Genes Expressed during Fruit Set Initiation in Tomato (Solanum Lycopersicum). PLOS ONE 2017, 12, e0180003, doi:10.1371/journal.pone.0180003.

30. Kim, D.; Langmead, B.; Salzberg, S.L. HISAT: A Fast Spliced Aligner with Low Memory Requirements. Nat Methods 2015, 12, 357–360, doi:10.1038/nmeth.3317.

31. Hosmani, P.S.; Flores-Gonzalez, M.; Geest, H. van de; Maumus, F.; Bakker, L.V.; Schijlen, E.; Haarst, J. van; Cordewener, J.; Sanchez-Perez, G.; Peters, S.; et al. An Improved de Novo Assembly and Annotation of the Tomato Reference Genome Using Single-Molecule Sequencing, Hi-C Proximity Ligation and Optical Maps 2019, 767764.

32. Vera Alvarez, R.; Pongor, L.S.; Mariño-Ramírez, L.; Landsman, D. TPMCalculator: One-Step Software to Quantify mRNA Abundance of Genomic Features. Bioinformatics 2019, 35, 1960–1962, doi:10.1093/bioinformatics/bty896.

33. Ge, S.X.; Son, E.W.; Yao, R. iDEP: An Integrated Web Application for Differential Expression and Pathway Analysis of RNA-Seq Data. BMC Bioinformatics 2018, 19, 534, doi:10.1186/s12859-018-2486-6.

34. Love, M.I.; Huber, W.; Anders, S. Moderated Estimation of Fold Change and Dispersion for RNA-Seq Data with DESeq2. Genome Biology 2014, 15, 550, doi:10.1186/s13059-014-0550-8.

35. Klavons, J.A.; Bennett, R.D. Determination of Methanol Using Alcohol Oxidase and Its Application to Methyl Ester Content of Pectins. J. Agric. Food Chem. 1986, 34, 597–599, doi:10.1021/jf00070a004.

36. Bradford, M.M. A Rapid and Sensitive Method for the Quantitation of Microgram Quantities of Protein Utilizing the Principle of Protein-Dye Binding. Analytical Biochemistry 1976, 72, 248–254, doi:10.1016/0003-2697(76)90527-3.

37. Bernard, C.; Traub, M.; Kunz, H.-H.; Hach, S.; Trentmann, O.; Möhlmann, T. Equilibrative Nucleoside Transporter 1 (ENT1) Is Critical for Pollen Germination and Vegetative Growth in Arabidopsis. Journal of Experimental Botany 2011, 62, 4627–4637, doi:10.1093/jxb/err183.

