## Supplemental Files for "Characterization and transcriptome analysis reveal abnormal pollen germination in cytoplasmic male sterile tomato"

**Table S1.** DEG analysis for *PME* and *PMEI* genes

| GeneID | Gene | logFC | logCPM | PValue | FDR | DEG (\|log$_2$ FC \| > 1, FDR < 0.1) |
|---|---|---|---|---|---|---|
| Solyc12g099410 | *PME* | 0.68 | 12.6 | 0 | 0 | no |
| Solyc01g066420 | *PME* | 0.41 | 13.48 | 0 | 0 | no |
| Solyc06g084620 | *PME* | 0.5 | 13.98 | 0 | 0.01 | no |
| Solyc12g099230 | *PME* | 0.61 | 6.3 | 0 | 0.02 | no |
| Solyc01g068120 | *PME* | 0.42 | 12.91 | 0 | 0.02 | no |
| Solyc05g054360 | *PME* | 0.34 | 12.31 | 0 | 0.02 | no |
| Solyc05g052110 | *PME* | 0.44 | 10.11 | 0 | 0.08 | no |
| Solyc03g078090 | *PME* | 0.27 | 8.33 | 0.01 | 0.13 | no |
| Solyc01g099940 | *PME* | -0.29 | 11.09 | 0.03 | 0.42 | no |
| Solyc01g066410 | *PME* | -1.37 | 0.13 | 0.04 | 0.48 | no |
| Solyc03g083840 | *PME* | 1.5 | -0.68 | 0.13 | 1 | no |
| Solyc05g052120 | *PME* | 0.24 | 7.08 | 0.21 | 1 | no |
| Solyc01g066390 | *PME* | 1.65 | 8.18 | 0.23 | 1 | no |
| Solyc01g057220 | *PME* | -0.18 | 7.35 | 0.25 | 1 | no |
| Solyc11g005750 | *PME* | -0.2 | 3.88 | 0.34 | 1 | no |
| Solyc01g066360 | *PME* | 0.08 | 12.02 | 0.36 | 1 | no |
| Solyc03g078100 | *PME* | 0.54 | 0.38 | 0.42 | 1 | no |
| Solyc01g099950 | *PME* | -0.69 | -0.19 | 0.51 | 1 | no |
| Solyc07g017560 | *PME* | -0.06 | 11.43 | 0.54 | 1 | no |
| Solyc02g075620 | *PME* | -0.2 | 1.52 | 0.64 | 1 | no |
| Solyc01g099960 | *PME* | -0.97 | -1.15 | 0.67 | 1 | no |
| Solyc06g051960 | *PME* | -0.33 | -0.34 | 0.81 | 1 | no |
| Solyc11g005770 | *PME* | 0.25 | 0.81 | 0.82 | 1 | no |
| Solyc11g070187 | *PME* | 0.04 | 0.78 | 1 | 1 | no |
| Solyc11g070175 | *PME* | 0 | 2.2 | 1 | 1 | no |
| Solyc01g066425 | *PME* | 0 | -1.56 | 1 | 1 | no |
| Solyc01g067410 | *PME* | 0 | -1.56 | 1 | 1 | no |
| Solyc01g067420 | *PME* | 0 | -1.56 | 1 | 1 | no |
| Solyc01g079180 | *PME* | 0 | -1.56 | 1 | 1 | no |
| Solyc01g090130 | *PME* | 0 | -1.56 | 1 | 1 | no |
| Solyc01g091050 | *PME* | 0 | -1.56 | 1 | 1 | no |
| Solyc01g091060 | *PME* | 0 | -1.56 | 1 | 1 | no |
| Solyc01g098940 | *PME* | 0 | -1.56 | 1 | 1 | no |
| Solyc01g109740 | *PME* | 0 | -1.56 | 1 | 1 | no |
| Solyc02g014300 | *PME* | 0 | -1.56 | 1 | 1 | no |
| Solyc02g062150 | *PME* | 0 | -1.56 | 1 | 1 | no |
| Solyc02g080200 | *PME* | 0 | -1.56 | 1 | 1 | no |
| Solyc02g080220 | *PME* | 0 | -1.56 | 1 | 1 | no |
| Solyc02g081990 | *PME* | 0 | -1.56 | 1 | 1 | no |
| Solyc02g083830 | *PME* | 0 | -1.56 | 1 | 1 | no |
| Solyc03g083360 | *PME* | 0 | -1.56 | 1 | 1 | no |
| Solyc03g083870 | *PME* | 0 | -1.56 | 1 | 1 | no |
| Solyc03g123620 | *PME* | 0 | -1.56 | 1 | 1 | no |
| Solyc03g123630 | *PME* | 0 | -1.56 | 1 | 1 | no |
| Solyc04g079340 | *PME* | 0 | -1.56 | 1 | 1 | no |
| Solyc04g080530 | *PME* | 0 | -1.56 | 1 | 1 | no |
| Solyc05g047590 | *PME* | 0 | -1.56 | 1 | 1 | no |
| Solyc05g053680 | *PME* | 0 | -1.56 | 1 | 1 | no |
| Solyc06g009180 | *PME* | 0 | -1.56 | 1 | 1 | no |
| Solyc06g009190 | *PME* | 0 | -1.56 | 1 | 1 | no |
| Solyc06g034360 | *PME* | 0 | -1.56 | 1 | 1 | no |
| Solyc07g017600 | *PME* | 0 | -1.56 | 1 | 1 | no |
| Solyc07g043240 | *PME* | 0 | -1.56 | 1 | 1 | no |
| Solyc07g064170 | *PME* | 0 | -1.56 | 1 | 1 | no |
| Solyc07g064180 | *PME* | 0 | -1.56 | 1 | 1 | no |
| Solyc07g064190 | *PME* | 0 | -1.56 | 1 | 1 | no |
| Solyc07g065350 | *PME* | 0 | -1.56 | 1 | 1 | no |
| Solyc07g065360 | *PME* | 0 | -1.56 | 1 | 1 | no |
| Solyc08g078640 | *PME* | 0 | -1.56 | 1 | 1 | no |

| | | | | | | |
|---|---|---|---|---|---|---|
| Solyc09g059980 | *PME* | 0 | -1.56 | 1 | 1 | no |
| Solyc09g075330 | *PME* | 0 | -1.56 | 1 | 1 | no |
| Solyc09g075350 | *PME* | 0 | -1.56 | 1 | 1 | no |
| Solyc09g091730 | *PME* | 0 | -1.56 | 1 | 1 | no |
| Solyc10g049370 | *PME* | 0 | -1.56 | 1 | 1 | no |
| Solyc10g049380 | *PME* | 0 | -1.56 | 1 | 1 | no |
| Solyc10g049430 | *PME* | 0 | -1.56 | 1 | 1 | no |
| Solyc10g049440 | *PME* | 0 | -1.56 | 1 | 1 | no |
| Solyc10g049450 | *PME* | 0 | -1.56 | 1 | 1 | no |
| Solyc10g076430 | *PME* | 0 | -1.56 | 1 | 1 | no |
| Solyc10g077135 | *PME* | 0 | -1.56 | 1 | 1 | no |
| Solyc10g083820 | *PME* | 0 | -1.56 | 1 | 1 | no |
| Solyc11g050990 | *PME* | 0 | -1.56 | 1 | 1 | no |
| Solyc11g051010 | *PME* | 0 | -1.56 | 1 | 1 | no |
| Solyc11g051020 | *PME* | 0 | -1.56 | 1 | 1 | no |
| Solyc11g051030 | *PME* | 0 | -1.56 | 1 | 1 | no |
| Solyc12g008530 | *PME* | 0 | -1.56 | 1 | 1 | no |
| Solyc12g098340 | *PME* | 0 | -1.56 | 1 | 1 | no |
| Solyc09g092350 | *PMEI* | 1.24 | 10.22 | 0 | 0 | DEG |
| Solyc05g005030 | *PMEI* | 1.13 | 11.57 | 0 | 0 | DEG |
| Solyc02g069300 | *PMEI* | 1.23 | 11.49 | 0 | 0 | DEG |
| Solyc10g081670 | *PMEI* | 1.04 | 8.93 | 0 | 0 | DEG |
| Solyc01g088590 | *PMEI* | 0.67 | 2.15 | 0.03 | 0.35 | no |
| Solyc01g059940 | *PMEI* | -0.28 | 7.58 | 0.03 | 0.44 | no |
| Solyc03g112970 | *PMEI* | -0.36 | 1.34 | 0.41 | 1 | no |
| Solyc03g083700 | *PMEI* | -0.94 | -1.14 | 0.67 | 1 | no |
| Solyc03g083730 | *PMEI* | 2.03 | -1.44 | 1 | 1 | no |
| Solyc03g112990 | *PMEI* | 0.86 | -1.05 | 1 | 1 | no |
| Solyc01g088575 | *PMEI* | 0 | -1.56 | 1 | 1 | no |
| Solyc01g091100 | *PMEI* | 0 | -1.56 | 1 | 1 | no |
| Solyc01g091110 | *PMEI* | 0 | -1.56 | 1 | 1 | no |
| Solyc01g097660 | *PMEI* | 0 | -1.56 | 1 | 1 | no |
| Solyc02g069330 | *PMEI* | 0 | -1.56 | 1 | 1 | no |
| Solyc03g083660 | *PMEI* | 0 | -1.56 | 1 | 1 | no |
| Solyc03g083710 | *PMEI* | 0 | -1.56 | 1 | 1 | no |
| Solyc03g083720 | *PMEI* | 0 | -1.56 | 1 | 1 | no |
| Solyc03g083770 | *PMEI* | 0 | -1.56 | 1 | 1 | no |
| Solyc03g112170 | *PMEI* | 0 | -1.56 | 1 | 1 | no |
| Solyc03g112180 | *PMEI* | 0 | -1.56 | 1 | 1 | no |
| Solyc03g112960 | *PMEI* | 0 | -1.56 | 1 | 1 | no |
| Solyc04g015510 | *PMEI* | 0 | -1.56 | 1 | 1 | no |
| Solyc04g071850 | *PMEI* | 0 | -1.56 | 1 | 1 | no |
| Solyc06g005460 | *PMEI* | 0 | -1.56 | 1 | 1 | no |
| Solyc06g034370 | *PMEI* | 0 | -1.56 | 1 | 1 | no |
| Solyc07g042390 | *PMEI* | 0 | -1.56 | 1 | 1 | no |
| Solyc08g006690 | *PMEI* | 0 | -1.56 | 1 | 1 | no |
| Solyc08g007450 | *PMEI* | 0 | -1.56 | 1 | 1 | no |
| Solyc08g016100 | *PMEI* | 0 | -1.56 | 1 | 1 | no |
| Solyc08g016110 | *PMEI* | 0 | -1.56 | 1 | 1 | no |
| Solyc08g016120 | *PMEI* | 0 | -1.56 | 1 | 1 | no |
| Solyc08g077400 | *PMEI* | 0 | -1.56 | 1 | 1 | no |
| Solyc09g011280 | *PMEI* | 0 | -1.56 | 1 | 1 | no |
| Solyc09g011290 | *PMEI* | 0 | -1.56 | 1 | 1 | no |
| Solyc09g072950 | *PMEI* | 0 | -1.56 | 1 | 1 | no |
| Solyc09g082610 | *PMEI* | 0 | -1.56 | 1 | 1 | no |
| Solyc09g092170 | *PMEI* | 0 | -1.56 | 1 | 1 | no |
| Solyc10g018320 | *PMEI* | 0 | -1.56 | 1 | 1 | no |
| Solyc10g018330 | *PMEI* | 0 | -1.56 | 1 | 1 | no |
| Solyc10g076730 | *PMEI* | 0 | -1.56 | 1 | 1 | no |
| Solyc11g005470 | *PMEI* | 0 | -1.56 | 1 | 1 | no |
| Solyc11g005820 | *PMEI* | 0 | -1.56 | 1 | 1 | no |
| Solyc11g019910 | *PMEI* | 0 | -1.56 | 1 | 1 | no |
| Solyc12g009270 | *PMEI* | 0 | -1.56 | 1 | 1 | no |
| Solyc12g099190 | *PMEI* | 0 | -1.56 | 1 | 1 | no |
| Solyc12g099200 | *PMEI* | 0 | -1.56 | 1 | 1 | no |
| Solyc12g099210 | *PMEI* | 0 | -1.56 | 1 | 1 | no |

**Table S2.** RNA-Seq data used in this study

| Accession number | Cultivar | Tissue |
|---|---|---|
| DRR092919 | "Micro-Tom" | Root (1 week) |
| DRR092918 | "Micro-Tom" | Stem (1 week) |
| DRR092897 | "Micro-Tom" | Leaf (1 week) |
| DRR092895 | "Micro-Tom" | Leaf (3 weeks) |
| DRR092902 | "Micro-Tom" | Green Fruit |
| DRR092903 | "Micro-Tom" | Red Fruit |
| DRR092908 | "Micro-Tom" | Petal (0 DAF) |
| DRR092912 | "Micro-Tom" | Sepal (0 DAF) |
| DRR092904 | "Micro-Tom" | Pistil (0 DAF) |
| DRR092906 | "Micro-Tom" | Anther (3-4 mm bud) |
| DRR092905 | "Micro-Tom" | Anther (1 DBF) |
| DRR092907 | "Micro-Tom" | Anther (0 DAF) |
| PRJNA1250823 | "Micro-Tom" | 10 min-incubated pollen |

DAF; day after flowering

DAB; day befor flowering

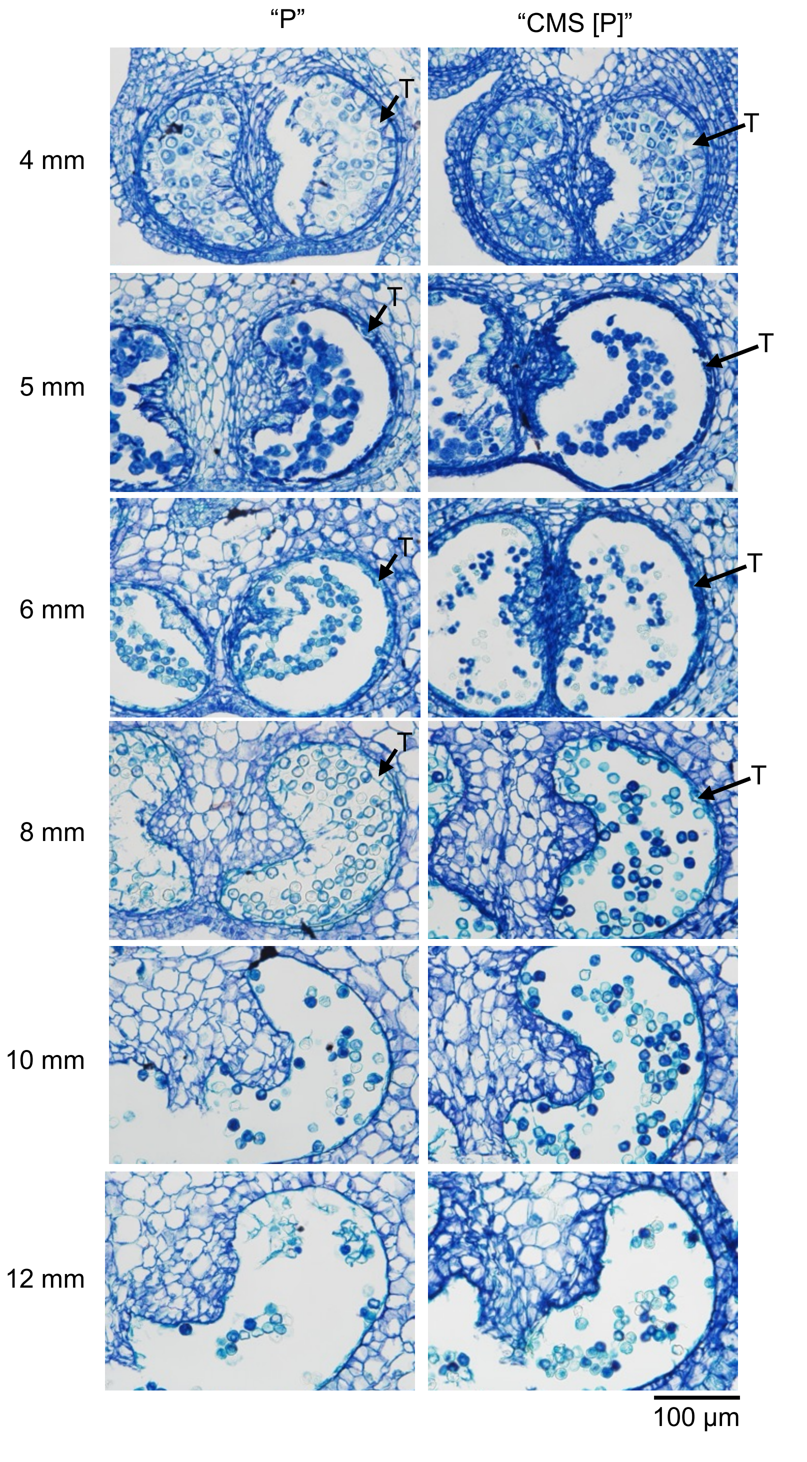


**Figure S1.** Developmental progression of anther tissues in "P" and "CMS[P]".
Cross-sections of anthers collected from flower buds of different sizes (4–12 mm) are shown.
T: tapetum layer.

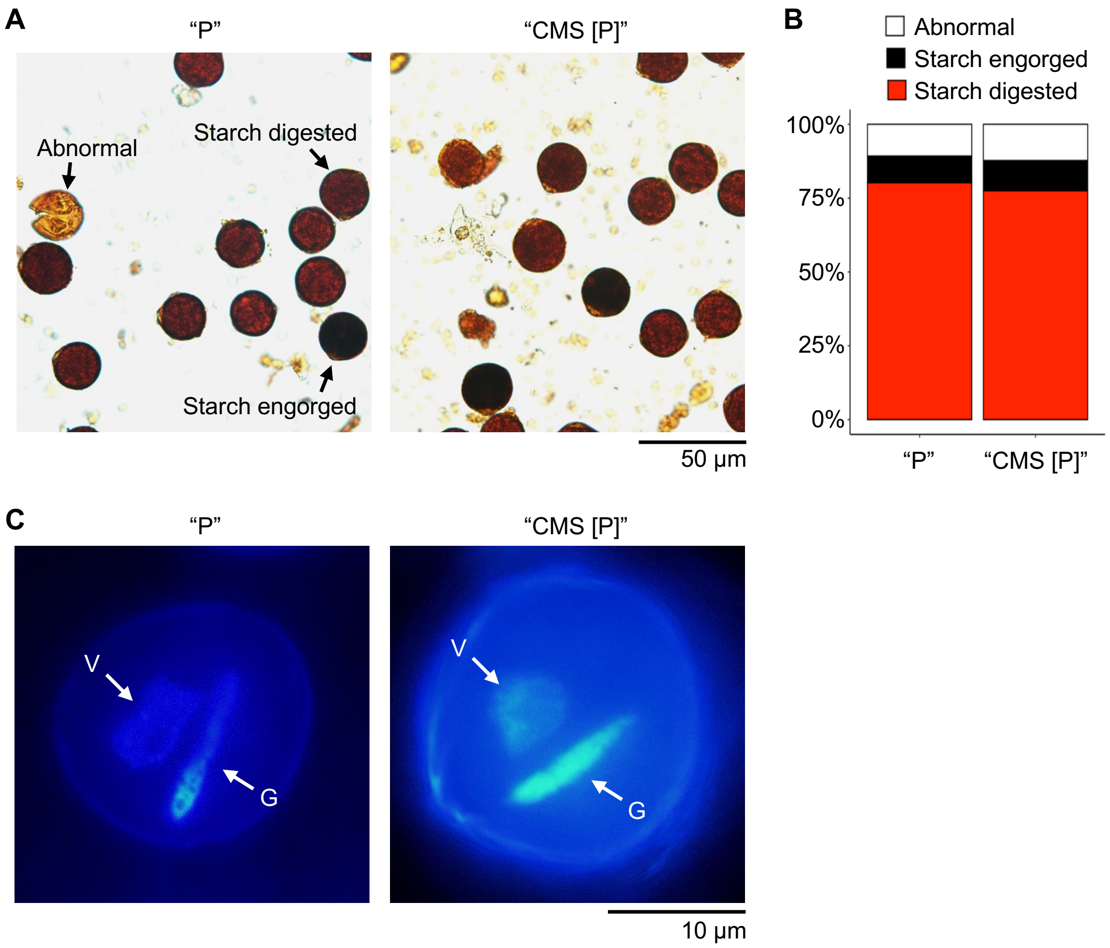


**Figure S2.** Pollen staining experiments in "P" and "CMS[P]".
(A) Pollen grains were stained by $I_2$-KI solution. Pollen was classified into 3 groups: starch digested, starch engorged, and abnormal pollen.
(B) Ratio of starch digested, starch engorged, and abnormal pollen in "P" and "CMS[P]". Data were obtained from five independent experiments (n = 5) and more than 500 pollen were counted in each experiment.
(C) Pollen grains were stained by DAPI solution. V; vegetative nucleus, G; generative nucleus.

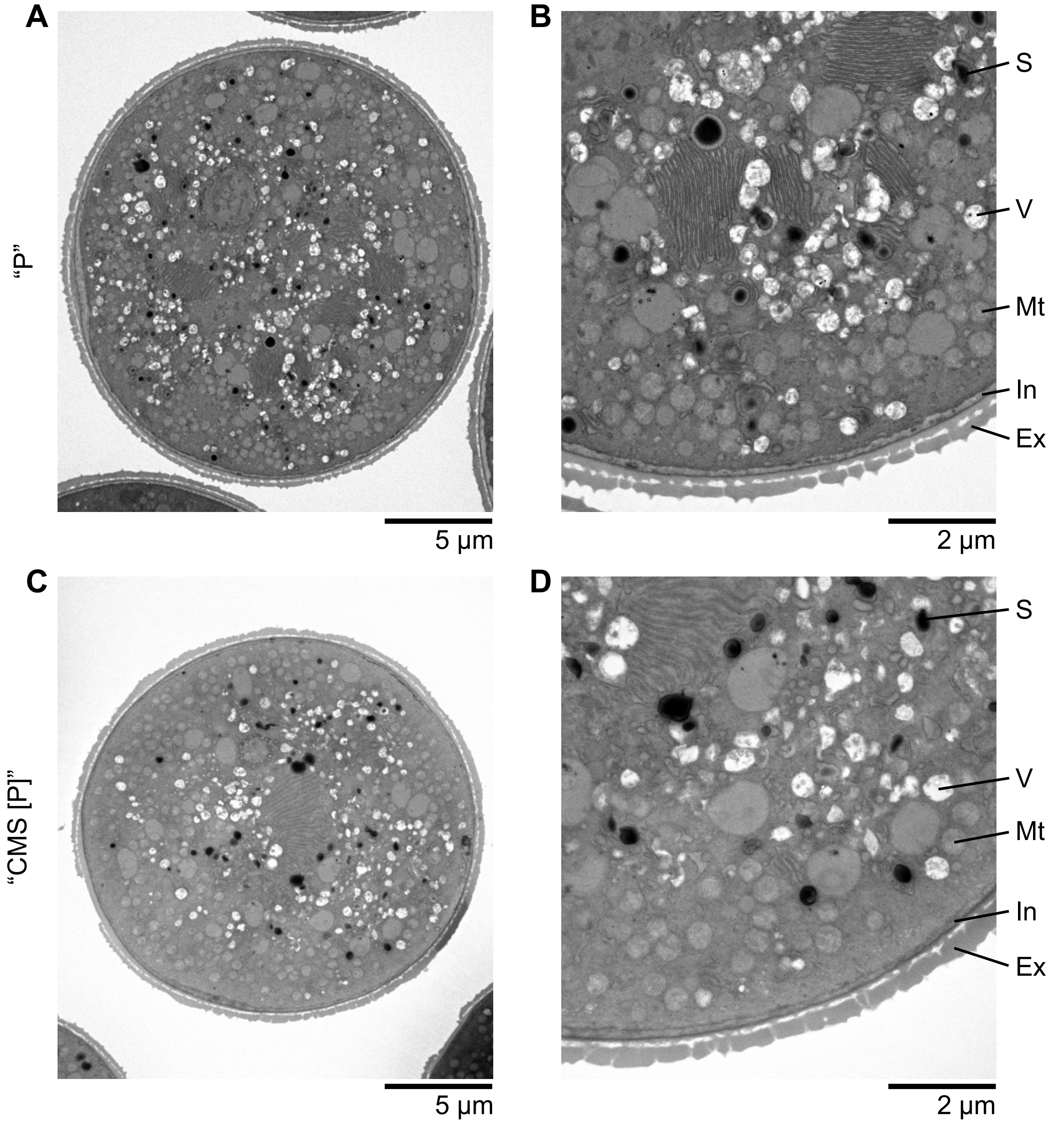


**Figure S3.** Transmission electron microscopy (TEM) analysis for pollen structure in "P" and "CMS[P]".

(A, B) TEM images of pollen structure in "P" and (C, D) "CMS[P]".

(B, D) Close-up pictures in A and C, respectively.

S; starch granule, V; vacuole, Mt; mitochondria, In; intine, Ex; exine.

**A** “Micro-Tom” Dwarf “CMS [P]”

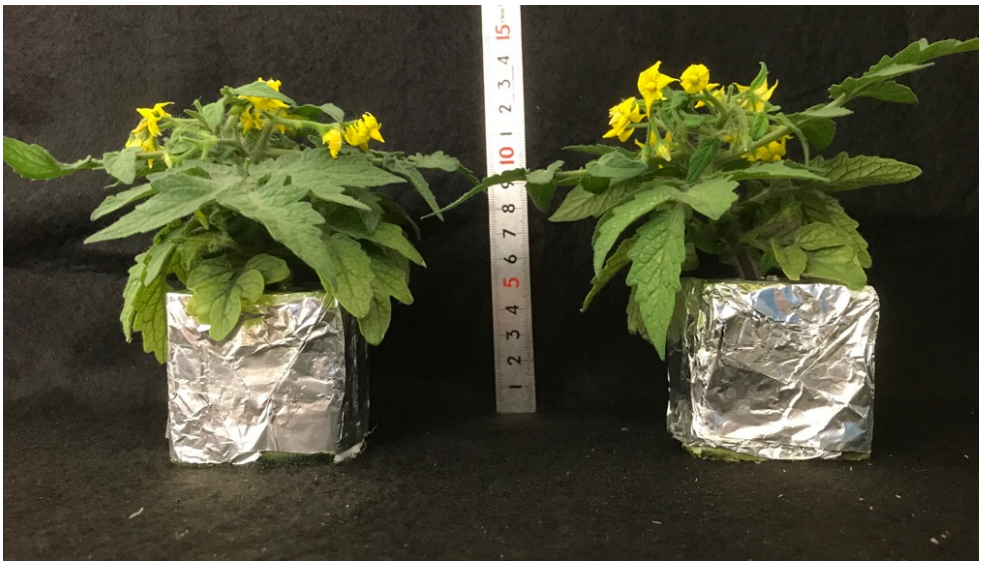

**B** “Micro-Tom” Dwarf “CMS [P]”

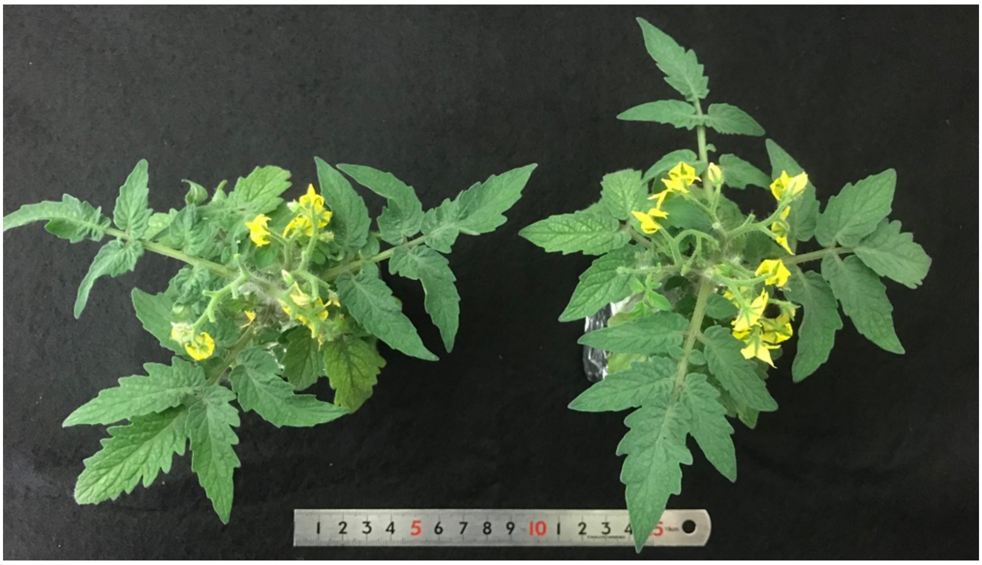

**Figure S4.** Plant morphology of “Micro-Tom” and Dwarf “CMS[P]”.
(A) Side view and (B) top view of flowering plants from “Micro-Tom” and Dwarf “CMS[P]”..
